# Real-World Plasma Proteomic Analysis of Pediatric Acute Recurrent and Chronic Pancreatitis

**DOI:** 10.64898/2026.11.24.768660

**Authors:** Dylan Nicholas Tabang, Oliver Schott, Zainab Wurie, Clara S. Gewinner, Xiaoyang Li, Fuchenchu Wang, Maisam Abu-El-Haija, Reuven Zev Cohen, Gretchen A Cress, Elissa M. Downs, Douglas S. Fishman, A. Jay Freeman, Gila Ginzburg, Tanja Gonska, Douglas Lindblad, Quin Liu, Asim Maqbool, Jacob Mark, Brian Arthur McFerron, Megha S. Mehta, Veronique D Morinville, Chee Y Ooi, Yuhua Zheng, Suyu Liu, Bernhard Y. Renard, Liang Li, Jami L. Saloman, Kinga K. Smolen, Mark E. Lowe, Aliye Uc, Hanno Steen, Chronic Pancreatitis Clinical Research Consortium (CPCRC, formerly CPDPC)

## Abstract

Pancreatitis is understudied in children compared to adults. Recent studies demonstrate the predominance of genetic risk factors over environmental factors in determining progression from acute to recurrent pancreatitis (ARP) attacks to chronic pancreatitis (CP) in children. Little has been done to describe the biomolecular profile in children developing CP. Plasma from 178 pediatric pancreatitis patients underwent mass spectrometry- and nucleic-acid linked immunosorbent assay-based proteomic analysis to distinguish ARP from CP, quantifying over 1000 proteins. Despite the unconventional sample handling and storage of these samples, whole blood collected in anti-coagulant, transported at ambient temperatures, and stored unfrozen up to ten days prior to plasma preparation, relevant pancreatitis biology was observed and a panel of five proteins distinguished ARP and CP with an AUC of 0.95 (95% CI 0.91-0.98). This analysis showed that pancreatitis patients grouped independently of genetic mutations, emphasizing the divergence of genotype and phenotype/endotype in pancreatitis. Proteomics identified a spectrum of early- and late disease-associated proteomic changes despite the clinical diagnosis, further underscoring the need for patient sub-stratification to identify potential therapeutic targets for pancreatitis in children.

**Highlights:**

- Plasma proteomics was performed on 178 samples from The INternational Study group of Pediatric Pancreatitis: In search for a cuRE (INSPPIRE) Consortium.
- Mass spectrometry and NULISA from EDTA whole blood transported at room temperature for up to ten days before processing into plasma is viable.
- A five-protein panel can distinguish between pediatric acute recurrent (ARP) and chronic pancreatitis (CP) with an AUC of 0.95 (95% CI 0.91-0.98).
- PRSS1 (trypsinogen) protein abundance is independent of PRSS1 gene mutation and pancreatitis severity, regardless of diagnosis.
- Inter-diagnosis heterogeneity reveals changes in immune response, RNA binding, cytoskeleton reorganization, and metabolism associated with progression toward CP.

## Introduction

The incidence of pediatric pancreatitis has increased in the last two decades, now affecting ∼1/10,000 children. Pancreatitis is less studied in children than in adults and is primarily driven by genetics, as opposed to environmental factors like alcohol consumption and tobacco usage in adults (1, 2). *The INternational Study group of Pediatric Pancreatitis: In search for a cuRE* (INSPPIRE) Consortium was founded to study pancreatitis in children with greater focus. It has enrolled 1175 patients, collecting biospecimens from approximately 900 of them, through the year 2026 from 28 clinical sites in the US, Canada, Israel, and Australia (3, 4). Patient enrollment began in 2012, following patients and accumulating metadata including disease stage progression and genetic sequencing for various risk factor genes. Progression from acute pancreatitis (AP) attacks to chronic pancreatitis (CP) in a subset of individuals and is often marked by recurrent attacks with periods of recovery, known as acute recurrent pancreatitis (ARP), that may ultimately lead to irreversible organ damage. In children, the median time for progression from AP to CP is 3.79 years (5), which is especially concerning given that CP is a significant risk factor for pancreatic cancer (6–8) and is associated with significant morbidity. CP diagnosis is based on fibrosis and calcifications, which are apparent by radiology, as well as exocrine and endocrine dysfunction (9, 10). Much of the previous work on this patient cohort has focused on identifying risk factors and disease burden in these children (11, 12), though much less is known about biomolecular processes that accompany disease progression. Progression from AP to ARP to CP is ultimately diagnosed by costly and potentially invasive imaging and pancreas function tests which are not necessarily available in primary or secondary healthcare centers. Plasma markers for pancreatitis progression would thus be valuable for diagnosis and for understanding the biological basis and disease pathology. Disease stratification would be helpful for determining treatment strategies, which currently focus on alleviating pain, regardless of disease stage diagnosis, or ultimately removal of the pancreas via total pancreatectomy islet auto transplantation (13–15).

Body fluid proteomics has proven useful in monitoring the overall picture of disease through progression or treatment in several conditions. Detection of proteins secreted into plasma through tissue leakage has also illuminated mechanisms in disease pathology. Plasma proteomics in pancreatitis in children has been under-explored, though a recent urine proteomics analysis of samples from a single center examined AP and CP versus healthy controls (16). Results from plasma proteomics, however, are described to be strongly affected by pre-analytical variation (17, 18). Prominent sources of variation include anti-coagulant choice, storage conditions, and whole blood centrifugation conditions. The presence of platelets, erythrocytes, and other blood cells is a major source of protein contamination which can inflate the number of species detected beyond secreted proteins in the “classical” plasma proteome. Such pre-analytical variations have emphasized the need for standard operating protocols (SOP) in specimen collection, processing, and handling for large studies. For example, serum is preferred for lipase and amylase tests to help diagnose pancreatitis (19). SOPs for plasma proteomic studies generally require plasma to be isolated and frozen within four hours of blood collection with anti-coagulant, generally ethylenediaminetetraacetic acid (EDTA) for plasma proteomics (20). The rigorous controls in SOPs for minimizing pre-analytical variation may hamper what analyses are possible using a given biospecimen. However, a recent study from our lab has shown that deviations from conventional biospecimen SOPs, like using freshly collected citrate plasma for platelet collection, do not necessarily preclude unraveling relevant biology (21).

The INSPPIRE-2 study collected whole blood, as genomic analysis of blood cells was the major rationale for collecting specimens in this way. Considering the relative stability of samples for genomic analysis and necessary equipment requirements for isolating plasma, blood samples collected in EDTA were shipped at ambient conditions, held at 4 °C for varying times before and after the shipment and with varying levels of movement, to a central location for centrifugation prior to freezing and subsequent proteomic analysis. This method of collection and transport is unique among other pancreatitis consortia biospecimens (22), which emphasized more conventional plasma collection. Despite the unconventional and nominally sub-optimal treatment of plasma for proteomic analysis, the aim of this study was to analyze whether these samples were still able to provide biological insight into pediatric pancreatitis.

## Experimental Procedures

### Reagents and Tools Table

| **Reagent/resource** | Reference/source | Identifier/catalog no. |
|---|---|---|
| **Chemicals, enzymes, and other reagents** |  |  |
| **Ethanol** | Fisher Scientific | A962-4 |
| **Water** | Fisher Scientific | W6500 |
| **Ammonium bicarbonate** | Millipore Sigma | A6141-500G |
| **Urea** | Millipore Sigma | U5378-1KG |
| **Dithiothreitol** | Millipore Sigma | 43815 |
| **Iodoacetamide** | Millipore Sigma | I1149 |
| **Sera-Mag Speedbead carboxylate-modified [E3] magnetic particles** | GE Healthcare | 65152105050250 |
| **Sera-Mag Speedbead carboxylate-modified [E7] magnetic particles** | GE Healthcare | 45152105050250 |
| **Trypsin** | Promega | V5111 |
| **Formic acid** | Millipore Sigma | F0507 |
| **Acetonitrile** | Fisher Scientific | A955-4 |
| **1-propanol** | Sigma-Aldrich | 34871-1L |
| **Software** |  |  |
| **DIA-NN** | https://github.com/vdemichev/diann | Version 1.8.1 |
| **R Studio** | https://docs.posit.co/ide/user/ | Version 2026.05.1 |
| **Others** |  |  |
| **4titude 96 Round Deep Well Storage Microplate** | Thomas Scientific | 1149R05 |
| **Eppendorf Deepwell Plates 96-well, 1000 µL** | Eppendorf | 951032107 |
| **Disposable Reagent Reservoirs, 100 mL** | Thermo Scientific | PI15075 |
| **Axygen Multiple Well Reagent Reservoir with 4-Channel Trough** | Corning | RES-MW4-HP |
| **MTC Bio 12 well reservoir** | Innovative Laboratory Products | RES16MC-12-N |
| **Adhesive PCR Plate Foils** | Thermo Scientific | AB0626 |
| **Thermomixer C** | Eppendorf | 5382000015 |
| **OT-2 automated liquid handler** | Opentrons | N/A |
| **OT-2 96 Tip Rack 300 µL** | Opentrons | 999-00009 |
| **OT-2 96 Tip Rack 20 µL** | Opentrons | 999-00007 |

### Cohort enrollment

Plasma samples (one sample per patient) analyzed were collected from patients enrolled in INSPPIRE-2. All patient centers obtained local IRB (or equivalent) ethical approval, with clinical sites in the US covered under MD Anderson Cancer Center IRB ID:2020-1057. Patients were diagnosed as ARP or CP under previously defined criteria. In short, ARP diagnosis required at least two distinct acute pancreatitis episodes with complete resolution of pain between diagnoses, while CP required one irreversible structural change in the pancreas as determined by imaging or histopathology: 1) once on any imaging – ductal calculi, dilated side branches, parenchymal calcifications; 2) persistent for >2 months on any imaging: ductal obstruction, stricture, dilation, or irregularities; 3) surgical or pancreatic biopsy specimen demonstrating histopathologic features compatible with CP (acinar atrophy, protein plugs, infiltration with lymphocytes, plasma cells, macrophages). Inclusion and exclusion criteria for enrollment were previously described (4). Clinical metadata is provided as **Supplementary Table 1**.

### Experimental Design and Statistical Rationale

Samples were randomized across two 96-well plates balancing ARP (n=90) and CP (n=88) diagnoses. The study was designed as pilot to establish feasibility for plasma proteomics, though post-hoc power analysis suggests that with power = 0.8 and α = 0.05, the minimum detectable effect size for a Wilcoxon rank sum test with FDR (Benjami-Hochberg corrected) < 0.05 is approximately 0.65-0.75. This corresponds to log_2_ fold change ∼ 1.5×.

### Clinical specimen processing

Blood was obtained via venipuncture in an EDTA tube, held at 4° C for up to 4 days, and shipped at room temperature, before receipt from various clinical sites across the United States, Canada, and Australia to the University of Iowa where they were stored for an additional up to 4 days at 4° C, up to 10 days total since collection, before processing into plasma. Blood vials were spun at 2360 ×g (3500 rpm) for 10 min in an Allegra X 30r centrifuge. Plasma was then frozen and shipped on dry ice to Boston Children’s Hospital for proteomics processing.

### Bottom-up proteomics sample preparation

Plasma samples (1 µL) in 96-well plates were diluted to 60 µL with 8 M urea in 50 mM ABC. 96-well plates containing the samples were thawed to room temperature from storage at −80 °C before sample preparation. Samples were reduced after addition of DTT to 0.01 M for 30 min with shaking at 800 rpm in a ThermoMixer (Eppendorf) and alkylated after addition of IAA to 0.02 M for 30 min in the dark. Alkylation was quenched with a further addition of DTT and incubation for 10 min with shaking at 800 rpm before proceeding.

An Opentrons OT-2 automated liquid handler was used to facilitate the rest of the SP3 (single-pot, solid-phase-enhanced sample preparation) on-beads digestion. Briefly, Sera-Mag SpeedBeads (1:10 protein to beads w/w) were dispensed in and washed with water twice. The reduced and alkylated plasma mixture was then applied to the beads. After addition of absolute ethanol to 80% v/v, samples were incubated for 10 min at 1000 rpm shaking. Sample plates were then put back into the automated liquid handler for washing with 80% ethanol thrice, each time after collection of the beads for 3 min with a magnetic module. Trypsin (1:25 enzyme to protein w/w) was then added in 100 µL 50 mM ABC. Digestion proceeded for 2 h at 37 C at 1000 rpm shaking.

Sample plates were centrifuged for 1 min at 3200 ×g prior to quenching digests with addition of FA to 2% v/v. Approximately 200 ng of peptides (25 µL of digest) were loaded onto Evotips for desalting prior to MS analysis. Tips were activated through washing with 0.1% FA in ACN, incubation in 1-propanol, and washing with 0.1% FA in water. Loaded samples were then further washed with 0.1% FA in water prior to LC-MS injection.

### Liquid chromatography-mass spectrometry proteomics and database searching

LC-MS/MS analysis was performed with an EvoSep One LC system coupled to a timsTOF Pro 2 (Bruker). Separation was done using a 60 sample per day (SPD) method, corresponding to a 21-min LC gradients, using a 8 cm length × 150 µm I.D. C18 column (Pepsep, Bruker).

DIA files were processed using DIA-NN (version 1.8.1). Raw files were searched using the same FASTA as the DDA files and against an in-house generated spectral library comprising thousands of plasma protein tryptic digest data files. Protein inference was against genes, neural network classifier was set to single-pass, quantification was done using a robust LC/high precision strategy, cross-run normalization was RT-dependent, library generation was done using smart profiling, and speed and RAM usage was set for optimal results.

### NULISA proteomics

10 µL of each plasma sample thawed, diluted, and centrifuged at 2,200 ×g for 10 min assayed with the NULISAseq Inflammation Panel 250. Immunocomplex formation with the paired set of oligo-conjugated antibodies, capture with oligo-dT beads, release, and capture with streptavidin beads and ligation were performed on the automated Alamar ARGO HT system. The library of DNA reporters containing unique target-specific molecular identifiers (TMI) and sample-specific molecular identifiers (SMI) was pooled, amplified by PCR, purified and sequenced on the Element Biosciences’ AVITI system. Target counts were normalized using internal controls and log2 transformed prior to statistical analysis.

### Statistical analysis

Raw protein intensities were derived from the protein group matrix output from DIA-NN. Dropped outlier samples were removed using protein count. Samples with protein counts outside of 3*standard deviation of the mean number of proteins across all samples were dropped from further analysis. Proteins detected in fewer than one-fourth of samples were removed from further analysis. Log2-transformed protein intensities were temporarily imputed using k-nearest neighbors prior to ComBat plate correction, with the pancreatitis diagnosis as a protected covariate. Imputed intensities were removed before statistical analysis, which was done using R Studio and various R packages (tidyverse, rstatix, ComplexHeatmap, pROC, eulerr, sva, pcaMethods, ggplot, impute). All reported significant p values < 0.05 are BH-corrected unless otherwise stated.

## Results

### Assessing proteome data quality and sample feasibility for plasma proteomics via MS and NULISA

We performed LC/MS analysis on 178 patient samples from the INSPPIRE-2 cohort collected in the United States, Canada, and Australia. Patients’ age ranged from 4-19 years, with 90 diagnosed with ARP and 88 CP. Other cohort statistics are detailed in **Table 1**. In addition to untargeted MS analysis, targeted proteomic analysis was performed on 171 samples via the NULISAseq Inflammation panel, which measures cytokines, chemokines, receptors, and other immune-modulatory proteins.

**Table 1:** Demographic details of the analyzed cohort.

|  | ARP | CP | Overall |
|  | (N=91) | (N=87) | (N=178) |
|---|---|---|---|
| Sex |  |  |  |
| Female | 52 (57.1%) | 54 (62.1%) | 106 (59.6%) |
| Male | 39 (42.9%) | 33 (37.9%) | 72 (40.4%) |
| Age |  |  |  |
| Mean (SD) | 12.2 (3.92) | 12.8 (3.61) | 12.5 (3.77) |
| Median [Min, Max] | 13.0 [4.00, 18.0] | 14.0 [4.00, 19.0] | 13.0 [4.00, 19.0] |
| Institution |  |  |  |
| Cedars-Sinai Medical Center | 1 (1.1%) | 3 (3.4%) | 4 (2.2%) |
| Children's Healthcare of Atlanta | 2 (2.2%) | 0 (0%) | 2 (1.1%) |
| Children's Hospital Colorado | 5 (5.5%) | 4 (4.6%) | 9 (5.1%) |
| Children's Hospital Los Angeles | 9 (9.9%) | 4 (4.6%) | 13 (7.3%) |
| Children's Hospital of Pittsburgh of UPMC | 11 (12.1%) | 4 (4.6%) | 15 (8.4%) |
| Children's Hospital of Wisconsin | 2 (2.2%) | 0 (0%) | 2 (1.1%) |
| Cincinnati Children's Hospital Medical Center | 6 (6.6%) | 14 (16.1%) | 20 (11.2%) |
| Montreal Children's Hospital | 3 (3.3%) | 0 (0%) | 3 (1.7%) |
| Nationwide Children's Hospital | 1 (1.1%) | 2 (2.3%) | 3 (1.7%) |
| Riley Children's Hospital at Indiana University | 5 (5.5%) | 8 (9.2%) | 13 (7.3%) |
| Stanford University School of Medicine | 4 (4.4%) | 6 (6.9%) | 10 (5.6%) |
| Texas Children's Hospital | 17 (18.7%) | 4 (4.6%) | 21 (11.8%) |
| The Children's Hospital of Philadelphia | 3 (3.3%) | 2 (2.3%) | 5 (2.8%) |
| The Hospital for Sick Children | 2 (2.2%) | 0 (0%) | 2 (1.1%) |
| UCSF Benioff Children's Hospital | 2 (2.2%) | 9 (10.3%) | 11 (6.2%) |
| University of Minnesota Masonic Children's Hospital | 2 (2.2%) | 15 (17.2%) | 17 (9.6%) |
| UTSW Children's Medical Center Dallas | 15 (16.5%) | 10 (11.5%) | 25 (14.0%) |
| Washington University School of Medicine | 1 (1.1%) | 1 (1.1%) | 2 (1.1%) |
| Sydney Children's Hospital - University of New South Wales | 0 (0%) | 1 (1.1%) | 1 (0.6%) |

Initial analysis of INSPPIRE-2 plasma samples first examined whether the unconventional and “sub-optimal” handling and transportation of the whole blood precluded successful proteomics. **Figure 1A** depicts the sub-cohort analyzed in this work. Diagnosis did not differ significantly by age and the ratio of females to males was similar between ARP and CP. **Figure 1B-C** depict the journey from clinical collection to plasma, in which patient whole blood samples from 19 clinical sites in the United States, Canada, and Australia were refrigerated for up to four days before shipping at ambient temperature to the University of Iowa for centrifugation into plasma and freezing, taking up to ten days in total. This extended storage and transport time, far beyond many those in many contemporary plasma proteomics papers, conflicts with conventional plasma proteomics studies, which emphasize the isolation and freezing of plasma within 4-6 hours of blood collection (23).

**Figure 1:**
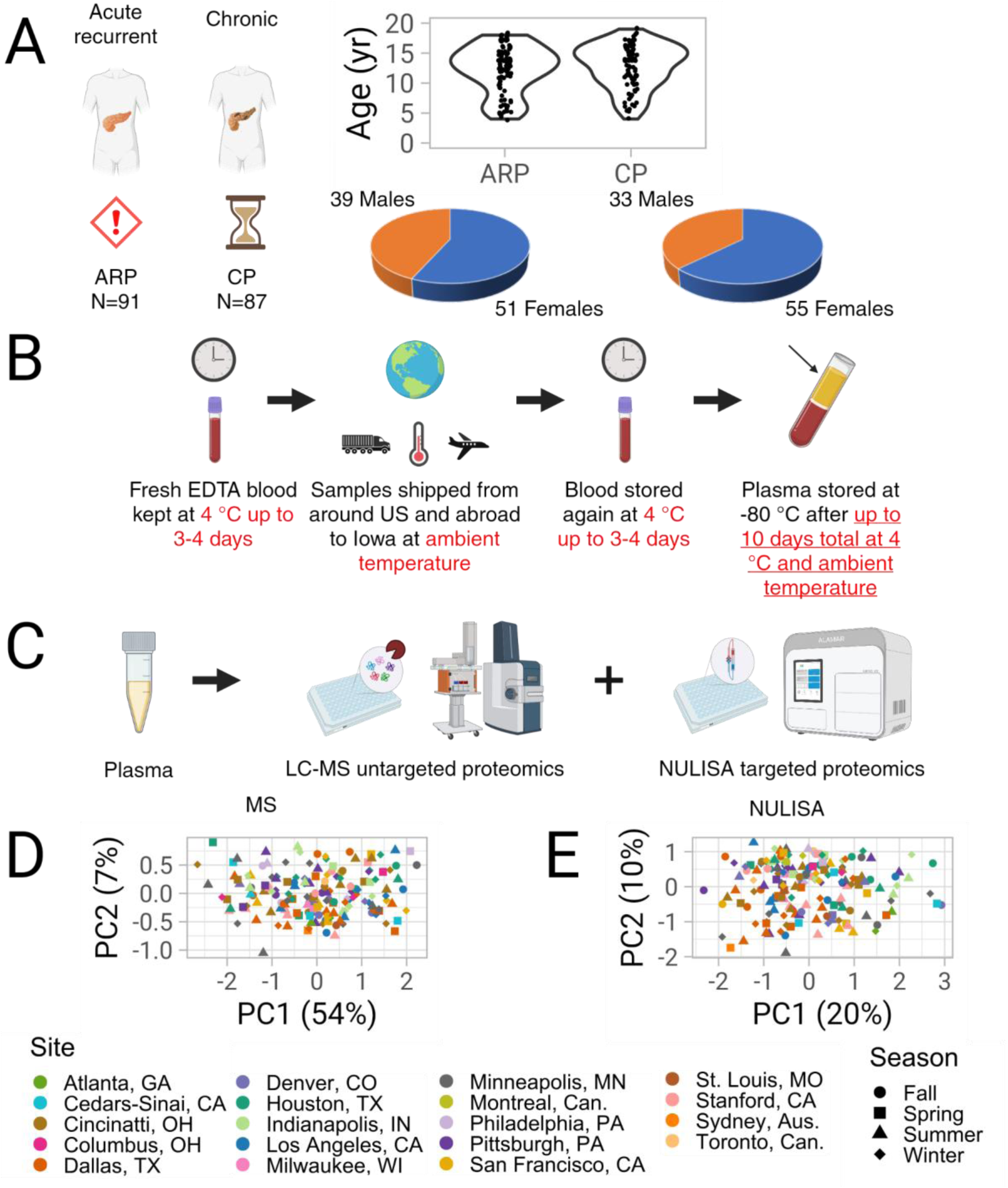
Workflow, cohort characteristics, and data overview. **A** Workflow of INSPPIRE blood samples from collection to plasma isolation. **B** Workflow of INSPPIRE plasma proteomics. **C** cohort pancreatitis, age, and sex ratio details. **D** PCA of all samples using mass spectrometry proteomics data showing that neither clinical collection site nor season the sample was drawn does not drive variation. The sample from Sydney, Australia is denoted with red arrow. **E** PCA of all samples using NULISA proteomics data.

Proteins were successfully extracted and digested into peptides for LC/MS analysis via diaPASEF on a Bruker timsTOF system, with spectra searched against an in-house generated spectral library comprising thousands of human plasma samples. This analysis identified 1110 unique proteins and on average 863 ± 71 (standard deviation, SD) proteins per sample. Hemolysis and secretion of proteins from other blood cells into plasma contributed to the higher detected protein count than expected. Three samples were excluded from further analysis due to low total protein count (<650 proteins, equivalent to ∼3 * standard deviation from the mean protein count). Only proteins detected in at least ¼ of all samples were kept for further analysis, totaling 1010 unique proteins (**Supplementary Figure 1A-B**). Batch correction to remove effects due to sample plate was performed using ComBat, with temporary imputation done using a k-nearest neighbors method.

MS intensities spanned ten orders of magnitude. The mean biological CV for proteins identified was ∼61%, though the median CV was only ∼48%. 541 out of 1010 quantified proteins (54%) had a CV < 50%. NULISA intensities also spanned ten orders of magnitude. The mean biological CV for the targets was ∼145%, with the median CV ∼86%. 51 out of 249 quantified proteins (20%) had a CV < 50%. This shift towards higher CV in the NULISA targets compared to MS is unsurprising, given the individual-specific variability in proteins related to immune signaling and other related pathways measured by NULISA and the potential impact of the suboptimal sample handling (**Supplementary Figure 1C-D**).

Examining the MS-detected proteins with CV <30% (170 proteins) reveals predominance of abundant proteins related to classical plasma pathways like complement activation and innate immune response. Example representative abundant proteins from classical plasma pathways include APOA1, F9, SERPINA4, and TF. An additional cluster of proteins in this group, including DDX5, HNRNPD, and TPR, reveals presence of proteins related to nuclear integrity and RNA processing. These pathways are also tightly controlled. Given the delayed processing of the samples and inevitable intracellular leakage, the presence of these proteins is not unexpected. Conversely, examination of MS-detected proteins with CV >100% (120 proteins) reveals the presence of erythrocyte and platelet markers, including ACTG1, FLNA, MYH9, and CA1 (18). In this more variable set of proteins, various intra-cellular pathways including “actin filament-based process” and “cell-cell adhesion” are among the most significant (**Supplementary Figure 2**). The degree of cellular damage resulting from pancreatitis may be one factor contributing to the higher biological CV, though the pre-analytical conditions of sample handling affecting individual protein heterogeneity cannot be ruled out. Indeed, known markers for erythrocytes (*e.g.* CA1, ALDOA, and PNP) and platelets (*e.g.* TLN1, ITGA2B, and TUBB4B) are distributed across the dynamic range of these samples, rather than most abundant (**Supplementary Figure 3**). The distribution of blood cell markers across the dynamic range, rather than clustered at the highest intensity end of the range, suggests that contamination may not be a major factor in influencing protein intensities.

Due to the varying distances of clinical site sample collection from plasma isolation at the University of Iowa and the varying ambient climate conditions for transport, PCA of all samples after outlier removal using the MS-derived protein intensities was performed to investigate these variables (**Figure 1D**). This analysis did not show distinct clustering by clinical site nor by temporal season during which samples were drawn. The one sample from Sydney, Australia, which would have by far the longest shipment time, was not clearly separated from the other samples. This suggests that variation in the samples is driven by something other than sample collection and transport.

The NULISA data recapitulated the PCA derived using MS proteomics (**Figure 1E**). Again, no clear separation of samples by clinical site or temporal season was found, though higher variance explained in PC1 compared to the MS data was found (54%, compared to 20%). This may be explained by the targeted nature of the assay leading to fewer quantified species and that these fewer targets are biased towards proteins with shorter lifetimes. Out of 249 targets in the panel, 246 were detected in over half of the samples. This high detectability is not surprising, given that the sensitivity of NULISA has been previously reported down to attomolar (10^-18^ M) concentrations (24). Indeed, Cohen et al. examined the effect of clinical sample handling on IL-6, IL-10, IFNγ, and IL-2 measurements in plasma and found that while whole blood storage at room temperature up to ten hours results in decreased levels, storage at 4 °C improves stability (20). They also found that “vigorous handling” of samples via pneumatic tube transport and up to three freeze-thaw cycles did not significantly affect measurements of these specific cytokines. Though the current work analyzes samples held at longer times at room temperature and 4 °C than in the work by Cohen et al. with similar freeze-thaw and transport conditions, the variability in transport conditions across clinical sites can help ensure that signals are indeed biological and not artifactual. Eighteen proteins were detectable in both MS and NULISA, spread throughout the MS dynamic range (**Supplementary Figure 4A**). Most targets were positively correlated between the two measurements, though only seven were both significant and had r > 0.5 (**Supplementary Figure 4B**). The strongest correlation (LCN2) and anticorrelation (S100A9) are shown in **Supplementary Figure 4C-D**.

### Proteomic analysis of recovery from AP attack

Having established that these samples generated robust and interpretable proteomic data, the next question was whether relevant pancreas biology was preserved in the samples. Nearly all samples analyzed (95%) had information on the time since last AP attack, which has not been previously explored longitudinally using human plasma samples. Patients were filtered into three groups based on the information of time between last AP attack and blood draw: <10 days (n=20), 11-40 days (n=20), and >1 year (n=55). Group sizes for the analysis of AP attack recency were determined based on known patient annotations, with a group size of at least twenty samples deemed sufficiently powered. A Kruskal-Wallis test identified 66 proteins significantly dysregulated among these three groups. A supervised clustering of patients using these proteins is shown in **Figure 2A**. The results show the distinct profile of patients who recently had an AP attack, who share an almost consistent proteomic response when compared to the convalescent patients in the heatmap. The two other groups have more heterogeneity, though the proteomic profile of patients whose last attack was approximately two weeks to a month before sample collection resembles the profile of those patients whose last attack was over a year before sample collection. This suggests that the plasma proteome “recovers” towards a pre-AP attack profile two weeks post-AP attack. **Figure 2B** shows the corresponding analysis in NULISA analysis, though the separation of three groups is less distinct than in the MS analysis. This is perhaps unsurprising, given the difference in average half-lives of cytokines/chemokines, *i.e.*, analytes measure by the inflammation panel, which is minutes to hours, compared to the bulk of the plasma proteome which have half-lives totaling days to weeks (25, 26).

**Figure 2.**
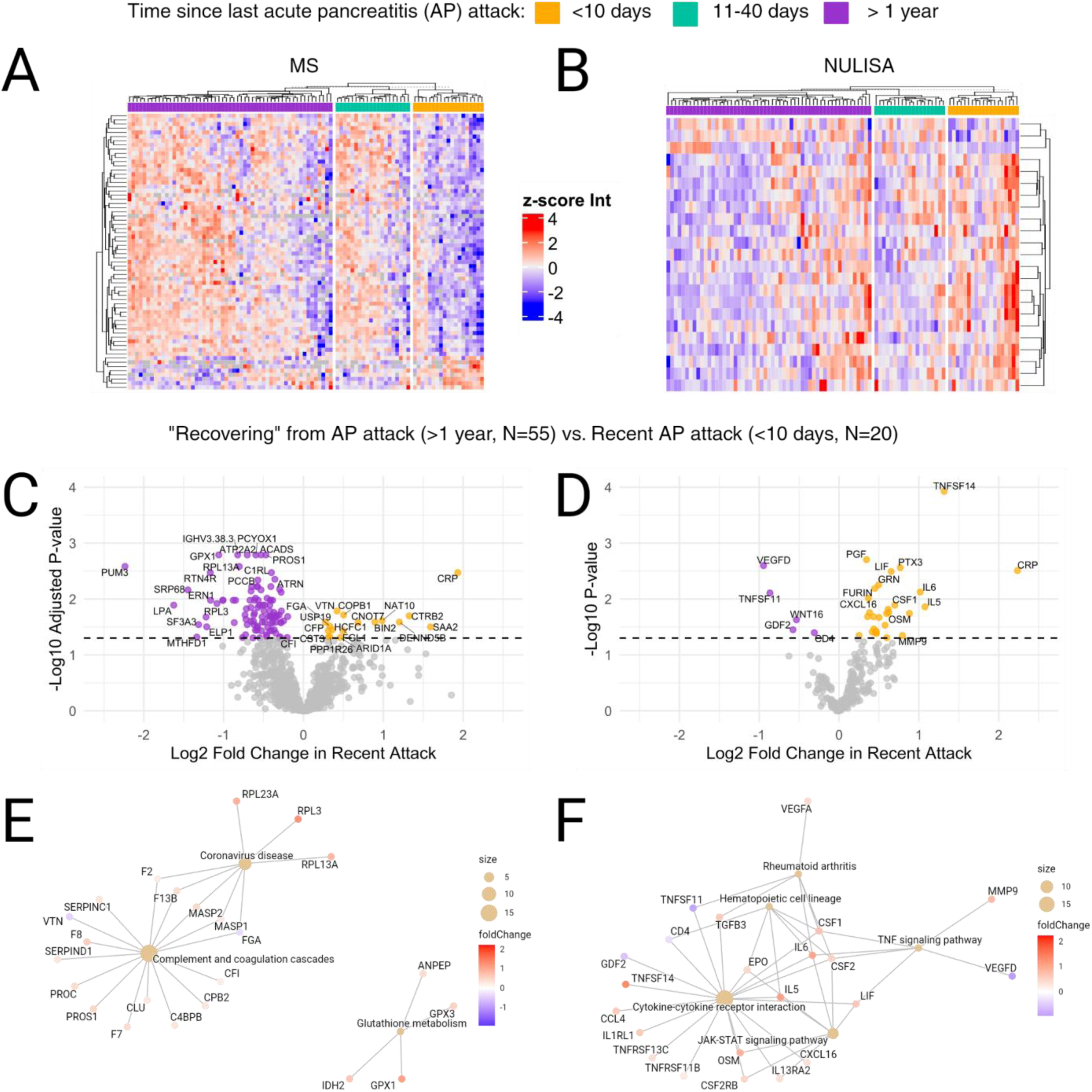
INSPPIRE blood sample feasibility for plasma proteomics is demonstrated through analysis of acute pancreatitis (AP) attack recency. **A** Supervised hierarchical clustering of patients using Kruskal Wallis BH-significant proteins detected by MS dysregulated throughout convalescence from an AP attack (<10 days, n = 20; 11-40 days, n=20; >1 year, n=55). **B** Supervised hierarchical clustering of patients using Kruskal Wallis significant proteins detected by NULISA. **C** Volcano plot of proteins detected by MS dysregulated in samples from patients who had an attack within the ten days prior to sample collection versus patients who have not experienced an AP attack in the prior year (Wilcoxon, BH-adjusted p <0.05). **D** Volcano plot of proteins detected by NULISA dysregulated in AP attack. **E** KEGG pathway enrichment analysis (p<0.05) of all BH-significant proteins detected by MS. **F** KEGG pathway enrichment analysis (p<0.05, top five pathways shown) of all BH-significant proteins detected by NULISA.

The binary comparison of patients with a recent attack (<10 days) and without a recent attack (>1 year, hereafter “recovering”) revealed further insights into AP attack biology, shown as volcano plots for MS and NULISA (nominal p <0.05 for NULISA) data, respectively, in **Figure 2C-D**. Importantly, age did not differ significantly between the two groups (Wilcoxon, p = 0.12). C-reactive protein (CRP), a known inflammatory marker, was upregulated in both MS and NULISA-based methods in recent attack. Other known inflammatory markers previously shown to be upregulated in AP, included SAA2, IL5, and IL6 (27, 28). Serum IL5 and IL6 were previously found to be associated with the severity of the AP attack (29). A previous antibody-based proteomics study in adult serum identified several proteins upregulated in pancreatitis (AP, CP, and recurrent AP combined) compared to healthy controls that were found here upregulated in patients with a recent attack, including OSM and TNFSF14 (30). This suggests a recovery in these proteins towards a more non-pancreatitis profile after one year without an AP attack. IL-17 and TNF signaling pathways were both upregulated in pancreas tissues of a rat model of AP compared to healthy control rats (31).

One major strength of MS proteomics is the untargeted nature of protein detection. MS detected upregulation of chymotrypsinogen B2 (CTRB2) in recent attack. This pancreas-secreted enzyme leaks into plasma due to the acinar cell injury results from the AP attack. Another digestive enzyme chymotrypsin-like elastase family member 2A (CELA2A) was similarly upregulated in recent attack but was filtered out due to detection in fewer than one-fourth of samples. This protein was previously found to be upregulated in urine from pediatric acute pancreatitis patients compared to healthy controls (16). It follows that CELA2A is upregulated in plasma in some patients who recently had an AP attack, as approximately half of total urine protein is derived from plasma (32). Further analysis of the MS data using Fisher’s exact test identified proteins whose observation frequencies significantly differed between recent and non-recent AP attack. PRSS2 and DCD were detected more frequently in recent the AP attack group, with PRSS2 being detected in 9/20 recent patients and 2/55 “recovering” patients (p=0.03) and DCD detected in 14/20 recent attack patients and10/55 “recovering” patients (p=0.03). PRSS2 is the less common isoform of trypsinogen, the inactive precursor of trypsin, while DCD is an innate host defense protein. PRSS2 was previously reported as a specific marker for AP through the urine trypsinogen-2 dipstick test, highlighted as a rapid and noninvasive diagnostic tool for pediatric AP specifically (33). While not significant (p=0.3), it is notable that CELA3B, another member of the elastase family, was detected in 8/20 recent attack patients versus only 5/55 “recovering” patients, again showing that digestive enzymes differentially leaking into the plasma could be readily detected despite the unconventional treatment of the samples.

**Figure 2E-F** highlights relevant pathways in the analysis of recent AP attack through KEGG enrichment of significant proteins and confirms detection of previously reported biology. JAK-STAT and TNF signaling pathways are highlighted in the NULISA data, though MS also highlights signatures related to glutathione metabolism in addition to complement and coagulation cascades. The processes associated with AP attack are consistent with the proteomic signatures identified here: 1) initial acinar cell injury leads to detection of PRSS2, CELA2A, and CTRB2 in plasma; 2) TNF signaling triggers inflammation that is spread by JAK-STAT signaling, as evident by upregulation of proteins including MMP9, IL6, and OSM; and finally 3) cellular changes detectable by MS, including the acute phase response (CRP, FGA), immune cell recruitment (BIN2, NAT10), complement activation (CFP, CST3), and other cellular damage leading to metabolic stress and transcriptional changes (USP19, COPB1). Enrichment of “glutathione metabolism” alludes to the resulting oxidative stress from the AP attack. Decreased GPX1 and GPX3 imply an inability to neutralize ROS, leading to mitochondrial injury. Decreased IDH2 further suggests further mitochondrial damage due to reduced NADPH and impaired recycling of GSH (34–36).

Based on the preserved previously reported biological signatures apparent from the patient samples, we concluded that questions about pancreatitis progression could be answered using these unconventionally handled plasma samples.

### Comparison of protein dysregulation in ARP versus CP

A central question of this work was whether plasma protein changes during ARP preceding irreversible organ damage defining CP could be detected. A Wilcoxon rank sum test identified 151 MS-detected proteins significantly dysregulated between the two diagnoses (**Figure 3A**). NULISA analysis identified 37 significant proteins, all upregulated in ARP (nominal p < 0.05). The NULISA panel used focuses on inflammation-related proteins, so it follows that significant proteins dominate the side of the volcano plot reflecting more acute inflammation in ARP over chronic inflammation in CP, the completion of disease progression in a persistently dysregulated state (**Figure 3B**). Supervised clustering of patients within the two diagnoses using these two sets of proteins highlights intra-diagnosis heterogeneity (**Figures 3C-D**). In the MS-detected proteins, CP patients cluster into one major group comprising 4/5 of patients. ARP patients, instead, cluster into three roughly equally sized groups, borders of which are denoted by arrows. In the NULISA-detected proteins, both diagnoses are quite heterogenous with less distinct clustering than with the MS-detected proteins.

**Figure 3.**
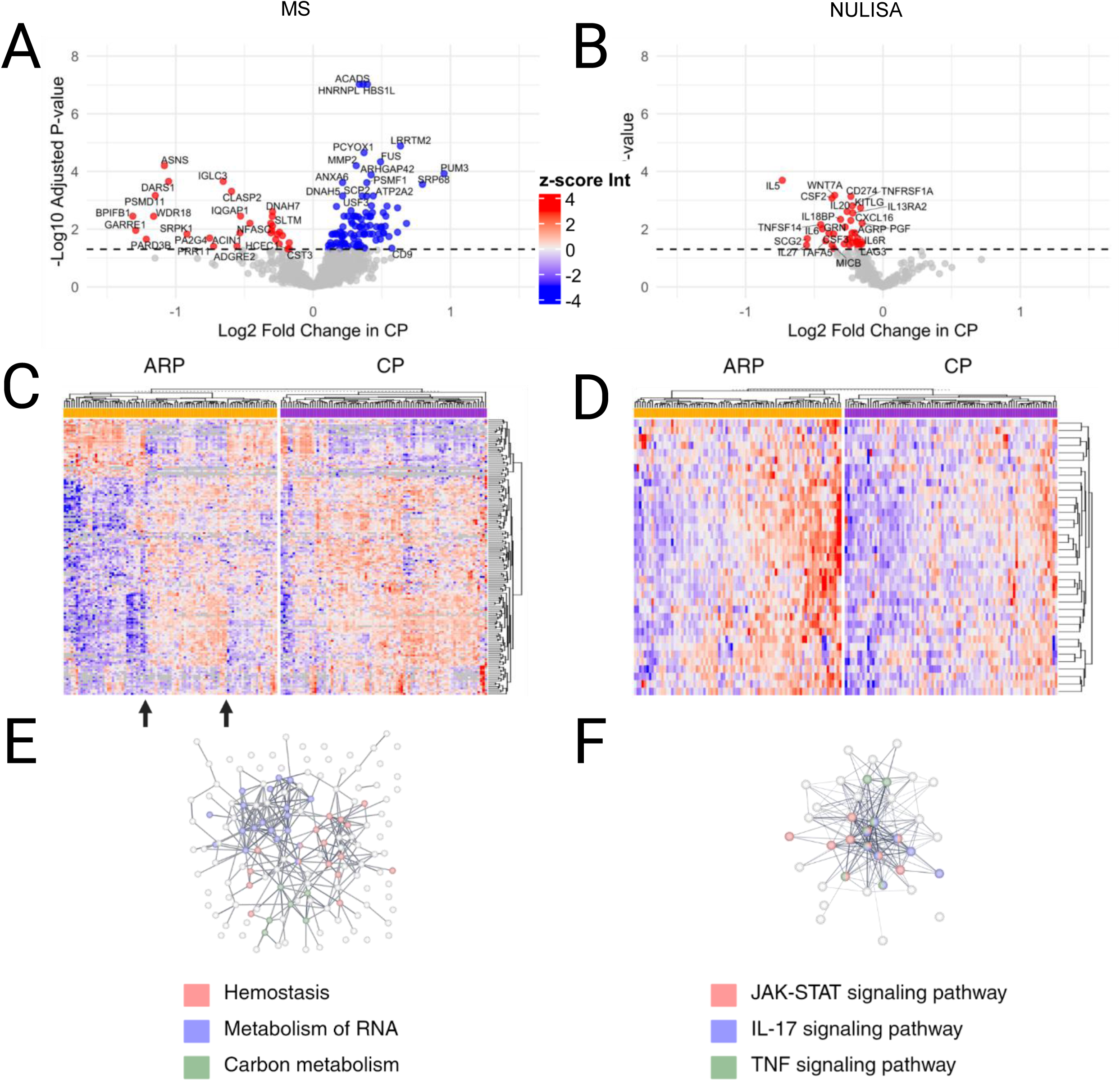
ARP vs. CP plasma proteomics reveals dysregulated proteins between diagnoses. **A** Volcano plot (Wilcoxon, BH-adjusted p <0.05) of proteins detected by MS dysregulated between ARP (n=89) and CP (n=86); **B** Volcano plot (Wilcoxon, nominal p <0.05) of proteins detected by NULISA dysregulated between ARP (n=84) and CP (n=86). **C** Supervised clustering of samples using MS ARP vs CP significant proteins reveals intra-diagnosis heterogeneity. **D** Supervised clustering of samples using NULISA ARP vs CP significant proteins similarly reveals intra-diagnosis heterogeneity**. E** STRING protein-protein interaction plot colored with pathway enrichment analysis (p<0.05) of all BH-significant proteins detected by MS. **F** STRING protein-protein interaction plot colored with pathway enrichment analysis (p<0.05) of all BH-significant proteins detected by NULISA.

STRING pathway enrichment analysis (**Figures 3E-F**) reveal similar terms as in the AP attack analysis but also reveals additional protein-protein interactions present in the proteins significantly different between ARP and CP, besides enriched pathways. The MS analysis reveals a major cluster of interacting proteins but also many proteins not interacting with others in the set. Three KEGG and Reactome pathway terms can explain a majority of the cluster: carbon metabolism, hemostasis, and metabolism of RNA. The NULISA analysis, which targets a set of known protein interactors, again highlights JAK-STAT and TNF signaling pathways, in addition to the IL-17 signaling pathway. IL-17 signaling is a known driver of pancreatitis progression and is also implicated in progression to pancreatic cancer (37, 38).

The MS analysis highlights upregulation of glutathione metabolism-related proteins in CP, as previously discussed, along with ACADS and PCCB, which are both involved in lipid metabolism. A previous analysis of hypertriglyceridemia (HTG) found that HTG severity was associated with increased pancreatic inflammation, and severe HTG had a significantly higher proportion of subjects with ARP versus CP (39). ACADS is an essential enzyme for mitochondrial fatty acid oxidation. The upregulation of ACADS in CP versus ARP may be reflective of a reduced oxidative capacity in ARP, leading to a lipotoxic environment promoting ongoing inflammation consistent with ARP.

### Deriving a protein panel for distinguishing ARP versus CP

**Figure 4** depicts the development of a panel of proteins distinguishing ARP vs. CP. Proteins were first filtered to a log_2_fold-change of >0.33, which corresponds to a ∼25% percent change between diagnoses, followed by an AUC filter > 0.7 for MS and > 0.6 for NULISA, which was set lower due to lower average AUC (**Figures 4A-B**). AUCs were calculated with age and sex as covariates. Every combination of up to five proteins was then examined for its AUC, sensitivity, and specificity without age and sex as covariates using logistic regression. For both MS and NULISA, an addition of up to five total proteins to the panel improved the AUC (**Figures 4C-D**). The best performing panel for MS, consisting of ARHGAP42, PSMD11, LRRTM2, HNRNPL, and PPP2R1A, distinguishing ARP and CP with an AUC from 0.91 to 0.98 (95% CI). The best performing AUC falls as the number of species in the panel decreases (0.80-0.91 for PSMD11, LRRTM2, HNRNPL and 0.71-0.84 for HNRNPL alone). For NULISA, the best performing panel, consisting of WNT7A, CSF3, IL17RB, IL6, and IL5, distinguished ARP and CP with an AUC from 0.83 to 0.93 (95% CI). As in MS, the best performing AUC range falls as the number of species in the panel decreases (0.67-0.82 for WNT7A, CSF3, IL17RB and 0.57-0.73 for WNT7A alone).

**Figure 4.**
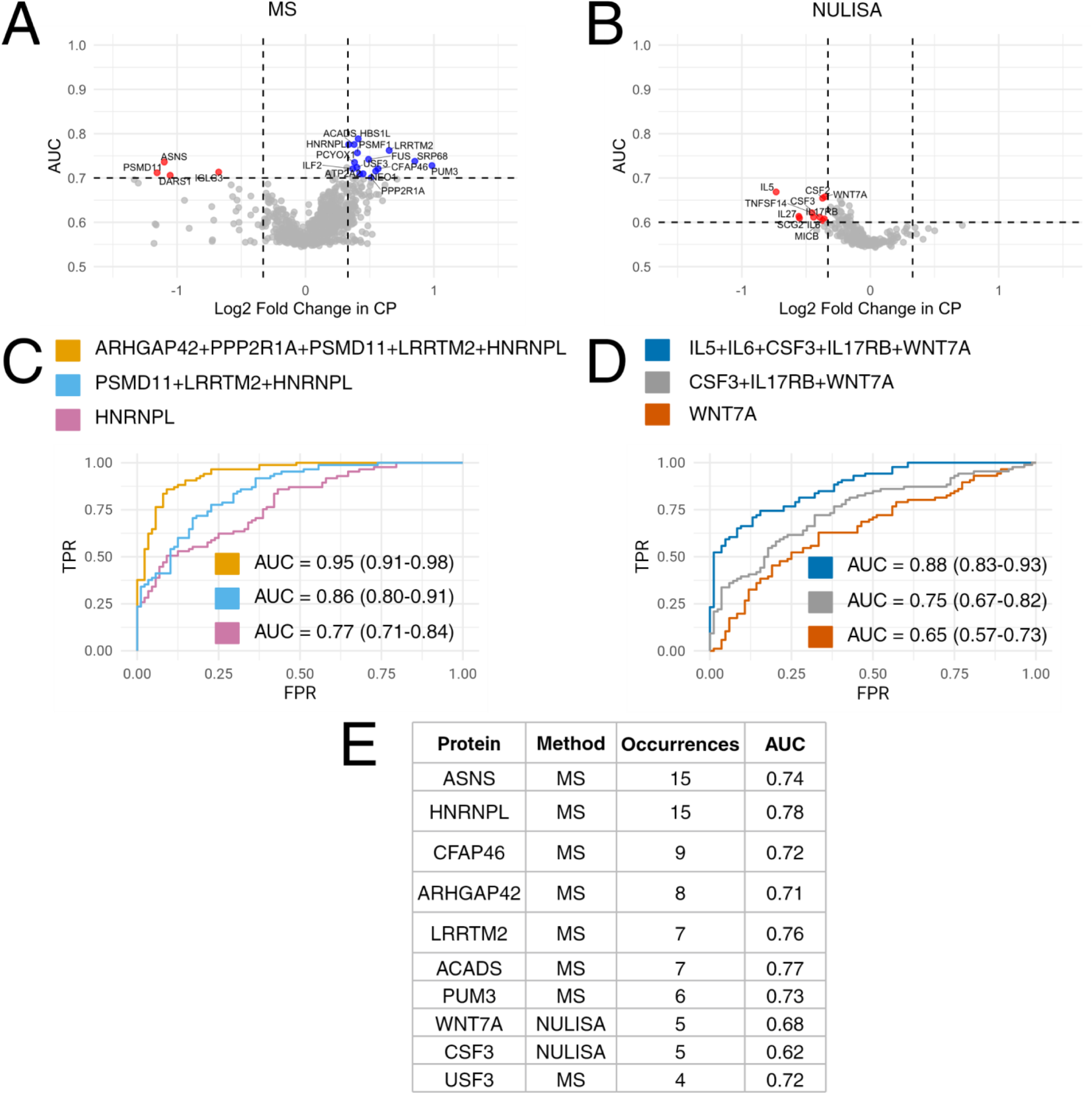
ARP vs. CP plasma proteomics reveals proteins that can discriminate between diagnoses. **A** Individual AUC values were calculated using MS detected proteins for CP vs ARP accounting for age and sex using a logistic regression model (GLM with bionomial family, model: Pancreatitis ∼ Intensity + Age + Sex. For future inclusion into protein panels for predicting ARP and CP, cutoffs of AUC > 0.7 and |log2 fold change| > 0.33 were used. **B** AUC values were similarly calculated using NULISA detected proteins, though a less strict cutoff of AUC > 0.6 was used with the same fold change cutoff as in MS. **C** A panel consisting of MS-detected proteins ARHGAP42, PSMD11, LRRTM2, HNRNPL, and PPP2R1A, was derived using all combinations of up to five proteins considering only proteins with a log2fold-change > 0.33 and AUC > 0.7, with the following curves based on highest AUC from three or one protein, respectively. AUC values and 95% CI for the three ROC curves. **D** A panel consisting of NULISA-detected proteins WNT7A, CSF3, IL7RB, IL6, and IL5, was derived using all combinations of up to five proteins considering only proteins with a log2fold-change > 0.33 and AUC > 0.6, with the following curves based on highest AUC from three or one protein, respectively. AUC values and 95% CI for the three ROC curves. **E** Table of the top ten proteins occurring in the top combinations for AUC in the combined MS+NULISA dataset, up to five proteins per panel.

IL6 has been repeatedly found as a plasma biomarker for CP compared to healthy controls (40). IL17RB is part of the IL-17 signaling pathway which in adult plasma distinguishes CP from AP and recurrent AP (30). The most significantly upregulated protein in ARP vs. CP is IL5. IL5 is being investigated as a therapeutic target for chronic muscle pain and fibromyalgia (41). Higher IL5 was found in patients, but exogenous IL5 in animal models was analgesic. ARP and CP differ significantly in their sampling times since last AP attack (Student’s t-test, p =0.0019), so higher IL5 in ARP may be indicative of ongoing inflammation. IL5 has been shown to promote progression of fibrosis and pancreatitis and is associated with AP severity (42), so patients with high IL5 may be more likely to transition to CP. Further longitudinal follow-up of INSPPIRE-2 patients will be necessary to test this.

To examine the robustness of single proteins in distinguishing ARP and CP, the AUCs of every combination of up to five proteins were calculated using logistic regression. **Figure 4E** shows a table of the top ten proteins in terms of number of occurrences in panels with AUC > 0.95, along with the AUC of the protein alone. ASNS and HNRNPL are likely robust markers for ARP vs. CP due to the much higher number of occurrences compared to the other proteins.

### Exploring effect of genetic risk factors and intra-diagnosis heterogeneity

The clustering of patients within diagnoses shown in **Figure 3C-D** emphasized patient heterogeneity, which we suspected may be due to their underlying genetic mutations causing their pancreatitis. Genetic mutations can explain up to 75% of pediatric pancreatitis cases (11, 43), with mutations in PRSS1, SPINK1, and CFTR being associated with most pediatric pancreatitis cases. The inactive precursor of trypsin, PRSS1, must first be cleaved to activate. Trypsin function is inhibited by SPINK1 or is flushed into the duodenum by CFTR. Premature activation of PRSS1 within the pancreas leads to autodigestion and inflammation which underscores pancreatitis. Patients, regardless of ARP or CP diagnosis, with known positive or negative status for mutation in each of these three genes were compared. **Figure 5A** shows volcano plots of these comparisons, with a gradient in number of significant proteins from left to right, from PRSS1 to SPINK1 to CFTR. This gradient mirrors the fact that PRSS1 mutation positive patients progress from ARP to CP faster than those with mutations in other genes, *i.e.*, ARP due to PRSS1 mutation is considered more severe (5, 44). SPINK1 and CFTR mutation positive patients see fewer significant proteins. This could be related to the more downstream effect of the mutation in the aberrant trypsin activation. While none of the proteins survived multiple testing correction, it is apparent that the PRSS1 mutation carriers are more dysregulated overall. Further data-driven exploration within each diagnosis was done considering the heterogeneity within ARP and CP respectively as previously shown in Figure 3C. Unsupervised clustering of patients within each diagnosis was first done using all proteins (**Figure 5B**). Based on the dendrogram, ARP or CP patients were divided into four or three groups, respectively. Mutation status in any of these three genes was not associated with the patient clustering in either diagnosis. Patients were separated into subgroups based on these clusters, with ARP having a cluster of patients with significantly higher PRSS1 expression (ARP3) than the other subgroups and CP with a subgroup of significantly lower PRSS1 expression (CP2, **Figure 5C**). This may correspond with time in disease progression where PRSS1 leakage from acini is highest during an AP attack and where PRSS1 expression has significantly decreased in CP due to excessive death of acini. This finding is noteworthy in that PRSS1 upregulation was not associated with mutation in the gene (**Figure 5D**). **Figures 5D-E** emphasize that when evaluating PRSS1 in known groups, mutation status (p=0.17) and pancreatitis diagnosis (p=0.52), respectively, PRSS1 expression does not significantly differ. This finding underscores that patient genotype does not dictate all aspects of endotype and overall plasma proteome expression.

**Figure 5.**
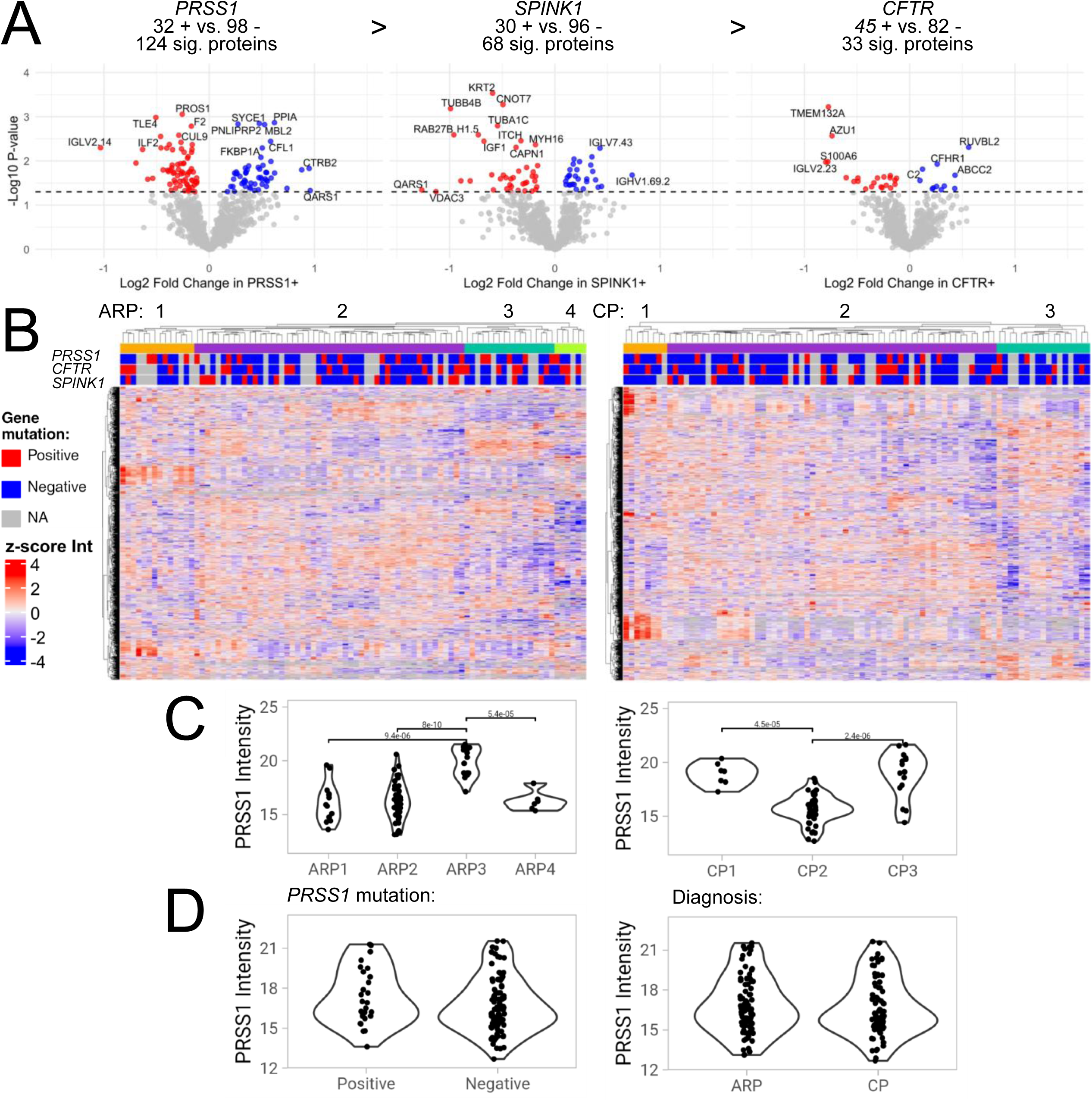
Proteomic analysis of common genetic risk factors for pancreatitis. **A** Volcano plots of proteins dysregulated in patients positive for a mutated gene versus those who are negative, for PRSS1, SPINK1, and CFTR. **B** Subgrouping of patients within ARP and CP diagnoses is independent of mutation status. **C** Subgrouping within ARP and CP diagnoses reveals upregulation of PRSS1 in a subpopulation without associated enrichment of PRSS1 mutation. **D-E** PRSS1 abundance does not significantly differ in comparisons of PRSS1 gene mutation status nor in ARP/CP diagnosis.

### Data-driven pan-diagnosis exploration of pancreatitis biology

The finding of PRSS1 dysregulation being independently driven raised the question of whether the progression of ARP to CP could be resolved using this dataset. The same patient subgrouping after unsupervised clustering methodology was repeated on all samples regardless of ARP or CP diagnosis to highlight disease-relevant pathways and identify disease progression trajectory. Patients were binned into six groups based on the dendrogram, with proteins filtered only to those that are significant after a Kruskal-Wallis test. Supervised clustering was then performed using 810 proteins with six groups of patients, which is depicted in **Figure 6A**.

**Figure 6.**
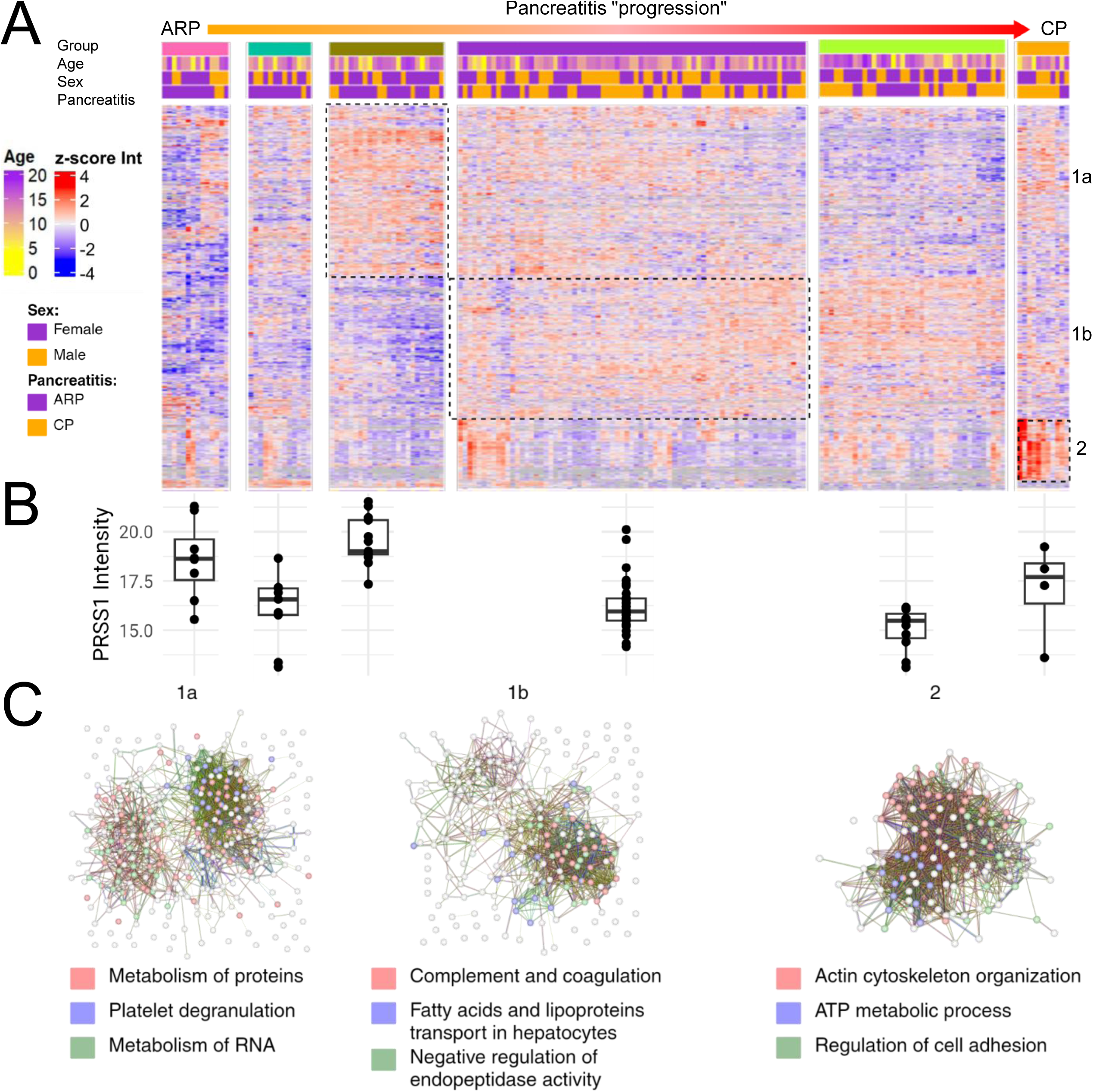
Subgrouping of pediatric pancreatitis patients reveals a progression from ARP to CP. **A** Supervised clustering of samples within subgroups using Kruskal Wallis significant proteins (BH-corrected p < 0.05) reveals a progression from ARP to CP, regardless of clinical diagnosis. **B** PRSS1 protein abundance varies among groups and throughout the ARP and CP progression. **C** STRING protein-protein interaction plots of clusters of proteins upregulated at three points in the ARP to CP progression.

Clustering was not associated with age, which did not differ significantly among the six groups (ANOVA, p = 0.72). Sex ratios within each group fluctuate between 1:1 and 2:1 female to male, though the olive group was 3:1. The orange group was dominated by CP, while pink, teal, and olive are mostly ARP. Differing ratios of CP:ARP, from 0.1 in the leftmost pink group to 5 in the rightmost orange group suggests possible disease progression depicted by group ordering. PRSS1 abundance is shown below each group in **Figure 6B**, which reveals a non-monotonic behavior for this protein across the six groups arranged from ARP to CP from left to right, though there is variability within each group. Variability is especially evident in the middle purple group, which is approximately 1:1 CP:ARP. Proteins were examined for enriched biological pathways using three separate STRING analyses (**Figure 6C**), with clusters 1a and 1b chosen for ARP-associated early pancreatitis changes and cluster 2 for CP-associated later pancreatitis changes.

AP begins with acinar cell damage from autolysis, which jumpstarts the body’s defense response, including complement activation, blood coagulation, and platelet degranulation. Negative regulation of proteolysis and endopeptidase activity is associated with the failure of these mechanisms leading to trypsin digesting the pancreas itself. One of the early disease-associated changes is RNA binding, which is a key part of the spliceosome. Dysregulation of the spliceosome has been linked to pancreatic cancer progression (45, 46). Fatty acid and lipoprotein transport may be disrupted due to AP attacks and ARP because of the spillage of lipase out of the pancreas. This contributes to the free fatty acid increase in systemic circulation, which is associated with lipotoxicity, AP severity, and eventually organ failure (47).

In the rightmost orange group, which is dominated by CP, the upregulated proteins in cluster 2 are enriched in terms related to the actin cytoskeleton and focal adhesion, as well as ATP metabolism. In both cerulein and diet-induced mouse models of pancreatitis, actin overexpression paralleled severity of pancreatic injury (48). Disruption of normal epithelial barrier integrity through actin dynamics during pancreatitis leads to leakage of cellular proteins which are then detected in the plasma (49, 50). Plasma peptidomics of a rat model of severe AP highlighted dysregulation of actin fragments (Acts, Actg), again emphasizing the importance of cytoskeletal remodeling during pancreatitis progression (51). Acinar ductal metaplasia, the transformation of acini into duct-like cells due to cellular injury from constant inflammation which involves reorganization of the cytoskeleton, may lead to precancerous lesions like PanIN, linking CP to pancreatic cancer (52).

Mitochondrial stress during progression from AP to ARP to CP leads to decreased ATP production. Although systemic hypercalcemia is an established driver of severe AP (53), the key cellular hallmark is sustained cytosolic calcium overload in acinar cells. The movement of Ca^2+^ from the endoplasmic reticulum (ER) and extracellular Ca^2+^ into the cytoplasm causes both toxic cytosolic calcium accumulation and ER calcium depletion. The resulting ER stress promotes the accumulation of unfolded proteins (54), further exacerbating mitochondrial dysfunction and feeding into the pro-inflammatory response.

Based on the different biological pathways enriched in the different groups of patients, this dataset appears to suggest a progression from ARP to CP. While protein panels could be used to differentiate ARP and CP, ultimately it is elucidation of the pathological mechanisms contributing to disease progression that will point towards targets for therapeutic intervention.

## Discussion

Overall, the work described here represents the first of its kind proteomics study examining plasma derived from blood collected in anti-coagulant and held at ambient temperature for extended periods. Despite this unconventional treatment of plasma leading to pre-analytical variation, previously reported biological features of acute pancreatitis attacks were seen and new insights into pediatric pancreatitis were derived. These findings highlight both proteomic differences between ARP and CP and enriched pathways throughout the progression of pancreatitis observed from proteomic profiles, which may conflict with the clinical diagnosis.

Improvements in mass spectrometry-based proteomics sensitivity and depth of coverage have enabled the analysis of large patient cohorts to derive insight into disease pathology. The diaPASEF-MS methodology employed here enabled the quantification of >800 proteins in approximately ∼20 min per sample from <1 µL plasma. The NULISA inflammation panel employed here quantified 250 protein targets, of which 18 could also be detected using MS, from 20 µL plasma. The unique proteins are of much lower abundance in plasma, thus requiring the more sensitive NULISA for detection. Though proteomics for biomarker discovery has improved in terms of depth and speed, barriers remain for these findings to be translated into clinical use, as discussed in a recent review (55). Particularly relevant to this work are the lack of harmonized reference frameworks, *i.e.*, standard baseline control samples, and pre-analytical variation of samples leading to misalignment between discovery experiments and clinical requirements. Various recent studies have examined the effect of pre-analytical variation on plasma proteomics, though all have concluded that secretion of cellular proteins into plasma may complicate the translational implications of the findings. The means of sample collection employed here highlight the main driver of pre-analytical variation, namely the presence of blood cells during ambient conditions and transport. This may have led to the higher variance in cytokines among all samples detected by NULISA due to the crosstalk between cell types during the stress of storage and transport. The putative biomarker proteins discussed here would need to be validated in samples treated in the same way that a clinical assay would be performed, *i.e.* plasma collected soon after blood collection.

Future studies into pancreatitis may benefit from the collection of dried blood spots (DBS), which are more stable in terms of sample transport and storage conditions (56). Collection of DBS does not require a phlebotomist and specialized equipment to isolate the plasma. This could prove useful for longitudinal collection, including pre- and post-AP attack. Urine has already proven a useful biospecimen for deriving insight into pediatric pancreatitis, with proteomics identifying biomarkers for the diagnosis of CP against controls and for fibrosis severity (57). The patient sub-stratification based on plasma proteome profiling described here requires the analysis of hundreds of patients to achieve appropriate statistical power. The non-invasive collection and relative stability of urine against rugged environmental conditions further make urine attractive as a sample matrix. Indeed, a recent large cohort study using urine explored genotype-proteome associations from >500 Parkinson’s disease patients (58).

The proteomics results recapitulated known AP attack biology, with the integration of NULISA and MS data revealing the interplay of cytokine surges and the cellular responses. The initial acinar cell damage from an AP attack directly led to the detection of digestive enzymes, *e.g.* CTRBS2, CELA2A, PRSS1, and PRSS2, in patients’ neat plasma. The initial cytokine surge, detectable in NULISA, spurred on signaling in the TNF, JAK-STAT, IL6, and IL17 pathways and leading to upregulation of proteins like TNFSF14, CSF1, and OSM in patients who recently had an AP attack but also in ARP compared to CP, which reflects the notion that ARP has more ongoing inflammation than CP. This is also supported by the information on time to closest AP attack from sample collection, which was significantly sooner in ARP patients compared to CP patients (280 days in ARP vs. 709 days in CP, p=0.002, **Supplementary Figure 5**). These changes in signaling co-occur with the cellular damage from leaking enzymes, creating a destructive feedback loop ultimately ending in permanent organ damage. Soon after initial acinar cell damage, the acute phase response (representative proteins include CRP, FGA), complement activation (CFP, PTX3), and immune recruitment occur (BIN2, MMP9).

Beyond the immune response which follows all cellular damage, stress to energy production, RNA processing, and protein folding machinery occur. Proteins in these pathways were useful in deriving a panel of proteins to distinguish ARP from CP. The best performing panel, with an AUC of 0.91-0.98 (95% CI) of MS-detected proteins consisted of ARHGAP42, PSMD11, LRRTM2, HNRNPL, and PPP2R1A. PSMD11 is a subunit of the 19S proteasome, a key part of the unfolded protein response. The buildup of unfolded proteins due to ER stress from the opening of calcium channels leads to upregulation of proteasomal subunits in AP. This ER stress leads to changes in the spliceosome, of which HNRNPL is a key mediator. With depletion of ATP from mitochondrial stress, cells switch towards necrosis from apoptosis, which is mediated by PPP2R1A. ARHGAP42 controls cell-cell adhesion and was upregulated in CP, which is associated with rearrangement of the cytoskeleton. Though LRRTM2 has not been previously described in plasma or pancreatitis contexts, its association with synapses can be linked to damage from leaking digestive enzymes spreading to sensory neurons around the pancreas (59, 60). Pancreatic neuropathy in CP has been associated with neural remodeling and neuropathic pain but LRRTM2 has not been investigated in this context (61).

Beyond the diagnostic implications of identifying proteins distinguishing ARP and CP, this study also enabled exploration of protein pathways changed throughout the progression from ARP to CP. Initial analysis was conducted on the most common genetic mutations associated with pediatric pancreatitis, specifically PRSS1, SPINK1, and CFTR. A comparison of patients based on mutation status, regardless of ARP or CP diagnosis, revealed a protein alteration gradient that correlates with disease severity. Specifically, PRSS1 gain-of-function mutations were associated with a stronger proteomic shift than other predisposing gene variants, such as SPINK1 and CFTR. Mutation status, however, did not dictate the similarity of patients within ARP or CP diagnoses, underscoring proteome heterogeneity and the importance of patient sub-stratification for future therapeutic considerations. This data-driven approach to clustering patients based on plasma proteome profile culminated in clustering of patients into six groups mirroring a progression from ARP to CP, though grouping was not necessarily by the clinical diagnosis of ARP to CP. The ARP-associated end of the grouping was associated with AP attack-related changes, like changes in protein and RNA metabolism and the immune response. The CP-associated end of the grouping was enriched in pathways related to cellular organization, like adhesion and cytoskeleton organization. The heterogeneity of patients in the middle groups requires further exploration but will also require more patients and higher statistical power to ensure confidence in the identified pathways.

Though this study was unique in its conception and methodology, there are some limitations to interpretation of the findings. One limitation is the variable transport and storage time of samples among the clinical sites. Stability of proteins in these conditions, *i.e.*, prolonged transport and storage at ambient temperatures exceeding one week, has not been explored previously, though it can be assumed that proteins do not have equal degradation and turnover rates. Though the biomarker panels derived here can distinguish ARP vs. CP, these findings would have to be validated in fresh plasma to not have to recapitulate the collection conditions here, *i.e.*, holding whole blood for an extended period before measurement, for any potential antibody-based assay. This study was done using ∼15% of available INSPPIRE-2 plasma biospecimens and these findings need to be validated. Conventionally derived plasma, isolated within four hours of whole blood collection, was not collected for this cohort. Transport and storage controls for these samples are thus not available for comparison. Another limitation affects the subgrouping analyses, both intra- and pan-diagnosis, which creates smaller groups by design. Though this study was statistically powered to investigate ARP vs CP, the small subgroup sizes make generalization of the findings more difficult by lowering the statistical power. Utilizing the full INSPPIRE cohort can illuminate whether biological signatures of patient subgroups are preserved. An additional limitation is not having pre-AP attack control patients, which would reveal changes in the plasma proteome in the first few days immediately post-attack.

In conclusion, we performed a proteomic analysis in pediatric pancreatitis plasma which not only recapitulated known biology of AP attacks, but also revealed differences between ARP and CP. This analysis of plasma derived from whole blood that underwent unconventional storage and transport conditions reveals that pristine protocols may not necessarily be required to derive biological insights into disease, assuming samples are treated to similar rugged storage and transport conditions. Our study shows that even blood samples handled “suboptimally” appear useful for plasma proteomics studies, expanding proteomics to clinical studies in less resourced settings. The knowledge gained about progression of pancreatitis in children can thus be utilized toward research into new treatments beyond pain management.

## Acknowledgments

All figures were generated in part using Biorender.

Research reported in this publication was supported by National Institute of Diabetes and Digestive and Kidney Diseases of the National Institutes of Health under award number(s) related to The Consortium for the Study of Chronic Pancreatitis, Diabetes, and Pancreatic Cancer (CPDPC) U01DK108328 and participating clinical center(s) University of Iowa (INSPPIRE) U01DK108334.

We would like to acknowledge Alamar Biosciences’ Technology Access Program for their support for the NULISA experiments. We would also like to thank the patients and their families, without whom this work would not be possible.

We would like to acknowledge all members of the Chronic Pancreatitis Clinical Research Consortium (CPCRC), formerly known as the CPDPC. The authors would like to acknowledge the continuous support by Dr. Aynur Unalp-Arida as the program official for the CPDPC since its inception, as well as the past support of project scientist Dr. Jose Serrano (2015-2025) and project consultant Dr. Dana Anderson (2015-2025).

The content is solely the responsibility of the authors and does not necessarily represent the official views of the National Institutes of Health.

## Data availability

LC-MS and NULISA protein data matrices are provided as **Supplementary Tables 2-3**.

The mass spectrometry proteomics raw files have been deposited to the ProteomeXchange Consortium via the PRIDE partner repository with the dataset identifier xxx (62).

## Author contributions

**Dylan Nicholas Tabang:** Conceptualization, Formal analysis, Investigation, Methodology, Visualization, Writing – original draft, Writing – review & editing. **Oliver Schott:** Formal analysis, Methodology, Visualization, Writing – review & editing. **Zainab Wurie:** Investigation, Writing – review & editing. **Clara S. Gewinner:** Investigation, Writing – review & editing. **Xiaoyang Li:** Data curation, Writing – review & editing. **Fuchenchu Wang:** Data curation, Writing – review & editing. **Maisam Abu-El-Haija:** Resources, Writing – review & editing. **Reuven Zev Cohen:** Resources, Writing – review & editing. **Gretchen A. Cress:** Data curation, Writing – review & editing. **Elissa M. Downs:** Resources, Writing – review & editing. **Douglas S. Fishman:** Resources, Writing – review & editing. **A. Jay Freeman:** Resources, Writing – review & editing. **Gila Ginzburg:** Resources, Writing – review & editing. **Tanja Gonska:** Resources, Writing – review & editing. **Douglas Lindblad:** Resources, Writing – review & editing. **Quin Liu:** Resources, Writing – review & editing. **Asim Maqbool:** Resources, Writing – review & editing. **Jacob Mark:** Resources, Writing – review & editing. **Brian Arthur McFerron:** Resources, Writing – review & editing. **Megha S. Mehta:** Resources, Writing – review & editing. **Veronique D. Morinville:** Resources, Writing – review & editing. **Chee Y. Ooi:** Resources, Writing – review & editing. **Yuhua Zheng:** Data curation, Writing – review & editing. **Suyu Liu:** Data curation, Writing – review & editing. **Bernhard Y. Renard:** Methodology, Writing – review & editing. **Liang Li:** Data curation, Writing – review & editing. **Jami L. Saloman:** Visualization, Writing – review & editing. **Kinga K Smolen:** Visualization, Writing – review & editing. **Mark E. Lowe:** Conceptualization, Funding acquisition, Supervision, Writing – review & editing. **Aliye Uc:** Conceptualization, Funding acquisition, Supervision, Writing – review & editing. **Hanno Steen:** Conceptualization, Methodology, Project administration, Supervision, Writing – original draft, Writing – review and editing, Writing – review & editing.

---

**Supplementary Table 1:** Other clinical metadata.

**Supplementary Table 2**: LC-MS data matrix.

**Supplementary Table 3**: NULISA data matrix.

**Supplementary Figure 1:** MS data quality metrics. **A** Violin plot of number of proteins identified in each sample. **B** Protein completeness curve depicting number of unique proteins identified in different fractions of the total samples analyzed. **C** Dynamic range of proteins identified in the MS and NULISA datasets, with rank inversely related to log-transformed intensity. **D** Density plots of the biological CVs of proteins identified in the MS and NULISA datasets.

**Supplementary Figure 2: A** Density of plot of biological CVs for proteins identified in the MS data split into two groups, <50% and >50%. **B** Gene ontology biological process enrichment (GOBP) of the two groups of proteins, sorted by –log(FDR). **C** STRING protein-protein interaction networks of the two groups of proteins.

**Supplementary Figure 3:** MS dynamic range annotated by presence of red blood cell and platelet marker proteins as highlighted by Geyer et al., 2019 (doi 10.15252/emmm.201910427).

**Supplementary Figure 4. A** Overlap of proteins detected by MS and NULISA and dynamic range of MS proteins annotated with the targets also detectable by NULISA. **B** Volcano plot depicting the Pearson correlation coefficients for the MS and NULISA intensities for the 18 shared targets. **C-D** Correlation plots for S100A9 and LCN2, which were the two targets with the most negative and most positive correlation coefficients.

**Supplementary Figure 5:** Boxplot depicting the days since last acute pancreatitis (AP) attack for patients split by ARP or CP diagnosis.

