## Supplemental Information for "Real-World Plasma Proteomic Analysis of Pediatric Acute Recurrent and Chronic Pancreatitis"

**Supplementary Information**

**Table S1**: Other clinical metadata.

**Table S2**: LC-MS data matrix.

**Table S3**: NULISA data matrix.

**Supplementary Figure 1:** MS data quality metrics. **A** Violin plot of number of proteins identified in each sample. **B** Protein completeness curve depicting number of unique proteins identified in different fractions of the total samples analyzed. **C** Dynamic range of proteins identified in the MS and NULISA datasets, with rank inversely related to log-transformed intensity. **D** Density plots of the biological CVs of proteins identified in the MS and NULISA datasets.

**Supplementary Figure 2: A** Density of plot of biological CVs for proteins identified in the MS data split into two groups, <50% and >50%. **B** Gene ontology biological process enrichment (GOBP) of the two groups of proteins, sorted by –log(FDR). **C** STRING protein-protein interaction networks of the two groups of proteins.

**Supplementary Figure 3:** MS dynamic range annotated by presence of red blood cell and platelet marker proteins as highlighted by Geyer et al., 2019 (doi 10.15252/emmm.201910427).

**Supplementary Figure 4. A** Overlap of proteins detected by MS and NULISA and dynamic range of MS proteins annotated with the targets also detectable by NULISA. **B** Volcano plot depicting the Pearson correlation coefficients for the MS and NULISA intensities for the 18 shared targets. **C-D** Correlation plots for S100A9 and LCN2, which were the two targets with the most negative and most positive correlation coefficients.

**Supplementary Figure 5:** Boxplot depicting the days since last acute pancreatitis (AP) attack for patients split by ARP or CP diagnosis.

**Supplementary Figure Legends**

**Supplementary Figure 1** MS data quality metrics. **A** Violin plot of number of proteins identified in each sample. **B** Protein completeness curve depicting number of unique proteins identified in different fractions of the total samples analyzed. **C** Dynamic range of proteins identified in the MS and NULISA datasets, with rank inversely related to log-transformed intensity. **D** Density plots of the biological CVs of proteins identified in the MS and NULISA datasets.

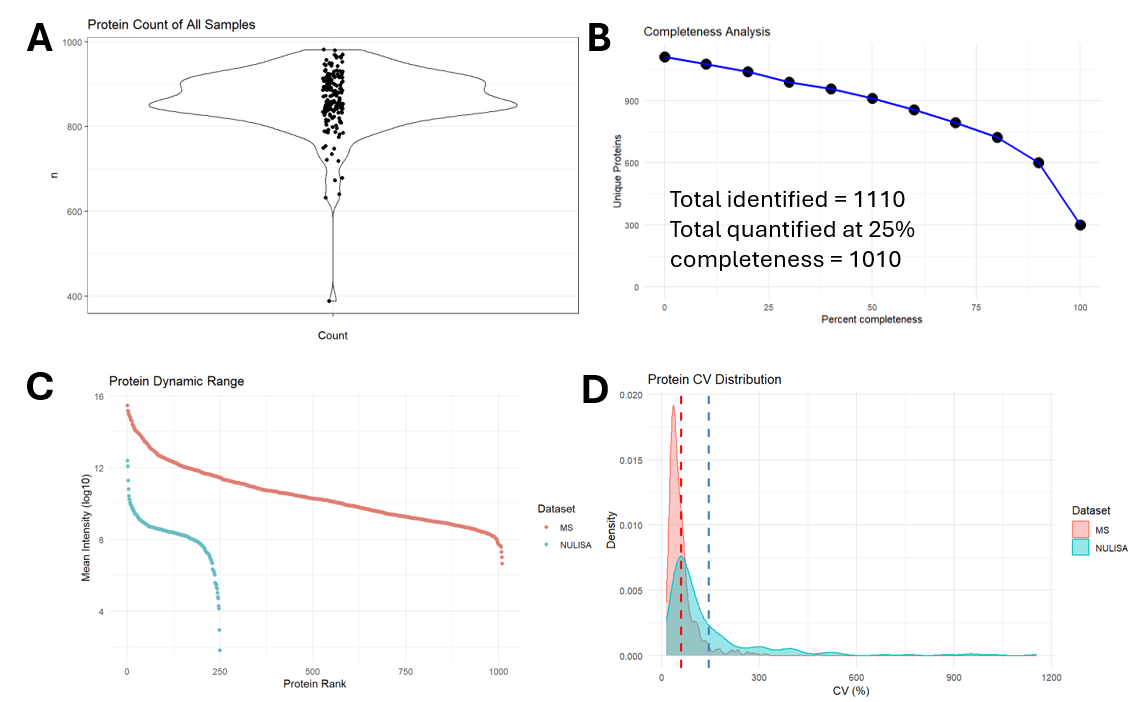

**Supplementary Figure 2 A** Density of plot of biological CVs for proteins identified in the MS data split into two groups, <50% and >50%. **B** Gene ontology biological process enrichment (GOBP) of the two groups of proteins, sorted by –log(FDR). **C** STRING protein-protein interaction networks of the two groups of proteins.
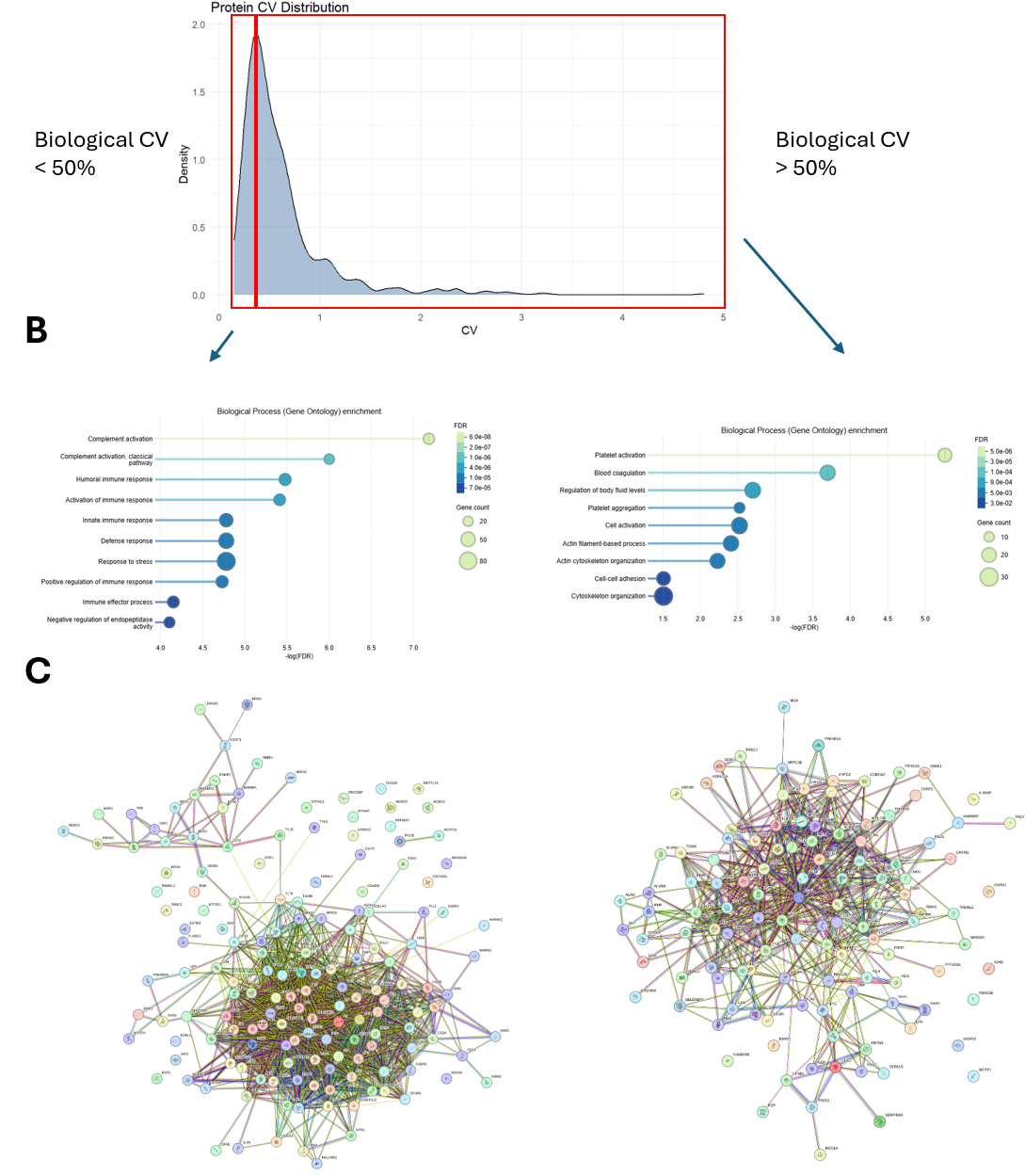

**Supplementary Figure 3** MS dynamic range annotated by presence of red blood cell and platelet marker proteins as highlighted by Geyer et al., 2019 (doi 10.15252/emmm.201910427).

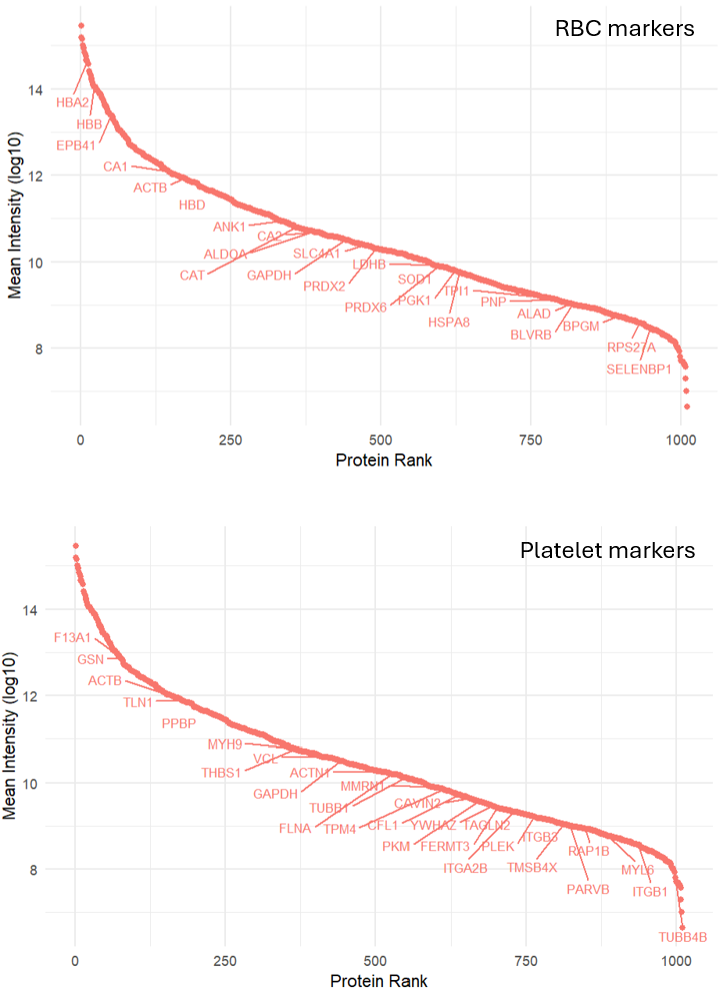

**Supplementary Figure 4 A** Overlap of proteins detected by MS and NULISA and dynamic range of MS proteins annotated with the targets also detectable by NULISA. **B** Volcano plot depicting the Pearson correlation coefficients for the MS and NULISA intensities for the 18 shared targets. **C-D** Correlation plots for S100A9 and LCN2, which were the two targets with the most negative and most positive correlation coefficients.

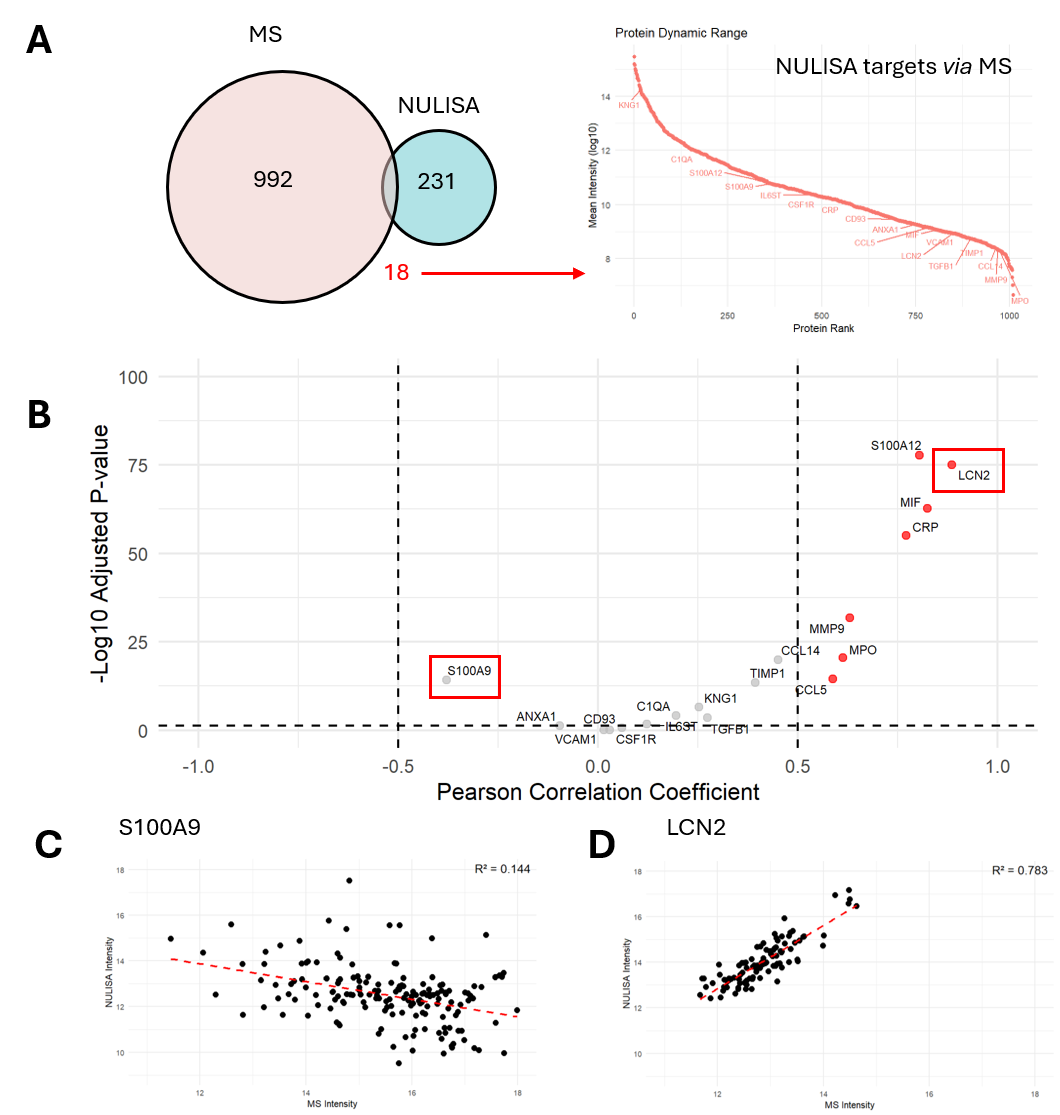

**Supplementary Figure 5** Boxplot depicting the days since last acute pancreatitis (AP) attack for patients split by ARP or CP diagnosis.

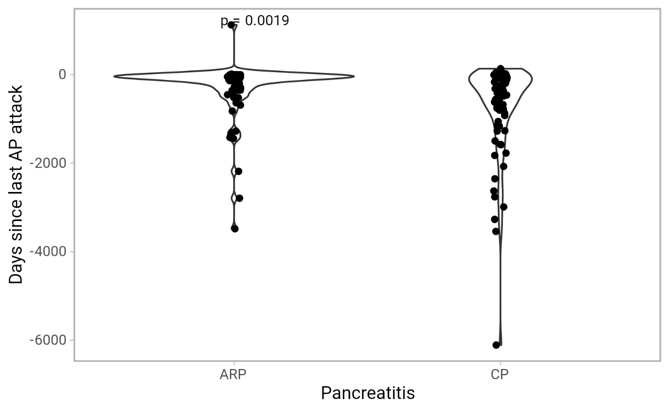
